# Experience directs the instability of neuronal tuning for critical period plasticity in mouse visual cortex

**DOI:** 10.1101/2025.01.15.633213

**Authors:** Thomas C. Brown, Aaron W. McGee

## Abstract

Brief monocular deprivation during a developmental critical period, but not thereafter, alters the receptive field properties (tuning) of neurons in visual cortex, but the characteristics of neural circuitry that permit this experience-dependent plasticity are largely unknown. We performed repeated calcium imaging at neuronal resolution to track the tuning properties of populations of excitatory layer 2/3 neurons in mouse visual cortex during or after the critical period, as well as in *nogo-66 receptor (ngr1)* mutant mice that sustain critical-period plasticity as adults. The instability of tuning for populations of neurons was greater in juvenile mice and adult *ngr1* mutant mice. We propose instability of neuronal tuning gates plasticity and is directed by experience to alter the tuning of neurons during the critical period.

## Main Text

Experience sculpts the connectivity and function of neurons in the central nervous system, particularly during developmental ‘critical periods’ when neural circuits are most plastic (Hensch & Quinlan 2018). This experience-dependent plasticity is best studied in the visual system where the tuning properties of neurons can be quantified from their responses to precise visual stimuli. In particular, the tuning for ocular dominance (OD) in visual cortex is conserved across numerous mammalian species including primates, felines, and rodents, and serves as a classic model for critical-period plasticity (Gordon & Stryker 1996; Hubel & Wiesel 1968; Huberman & Niell 2011; Priebe & McGee 2014; Wiesel & Hubel 1963). In the mouse, the critical period for OD plasticity spans postnatal day (P) 19 to ∼32 (Gordon & Stryker 1996). Closing one eye by lid suture (monocular deprivation, MD) for a few days only during this interval disrupts the normal binocularity by shifting the responses of neurons in primary visual cortex (V1) toward the open (non-deprived) eye (Gordon & Stryker 1996). Many genes, signaling pathways, and forms of synaptic plasticity, are required for OD plasticity (for review see (Levelt & Hübener 2012), but the features neuronal circuits that are permissive for this experience-dependent plasticity confined to the critical period remain poorly understood.

In recent work, we determined how OD plasticity alters the tuning of excitatory neurons in layer 2/3 of V1 by measuring their response properties both before and after MD (Brown & McGee 2023). MD evokes a complex reorganization of the tuning properties of neurons that shifts the overall binocularity of the population of visually-responsive neurons. Recent studies that have revealed a surprising instability of the tuning properties of neurons over time in multiple brain regions, a phenomenon termed ‘representational drift’ (Clopath et al. 2017). However, the physiologic role and significance of the phenomenon of representational drift remain unclear (Driscoll et al. 2022). Here we tested the hypothesis that greater instability of neuronal tuning is a defining circuit feature of plasticity during the critical period.

To determine the relationship between the instability of neuronal tuning in V1 and OD plasticity, first we confirmed that OD plasticity in response to brief MD is restricted to the critical period as measured with calcium imaging at neuronal resolution in alert wild-type (WT) mice with the sensor GCaMP6s (Fig. 1A-H, S1) (Brown & McGee 2023; Chen et al. 2013; Wekselblatt et al. 2016). MD during the critical period for 4 days alters the binocularity of the population of excitatory neurons in L2/3 and the average Ocular Dominance Index (ODI) per mouse decreased from values that reflect the typical bias of non-deprived mouse V1 for stimuli presented to the contralateral eye (Fig. 1 I-K) (Brown & McGee 2023; Frantz et al. 2020). This plasticity is confined to the critical period and is not detectable in adult mice (P60-90) as ODI values following 4 days of MD were similar to ND mice (Fig. 1I,H). These findings are consistent published studies that have examined the effects of brief MD in juvenile mice with multi-unit electrophysiologic recordings (Gordon & Stryker 1996), as well as more indirect methods such as visually-evoked potentials (VEPs) and optical imaging of intrinsic signals (Cang et al. 2005; Frenkel & Bear 2004).

**Fig 1.**
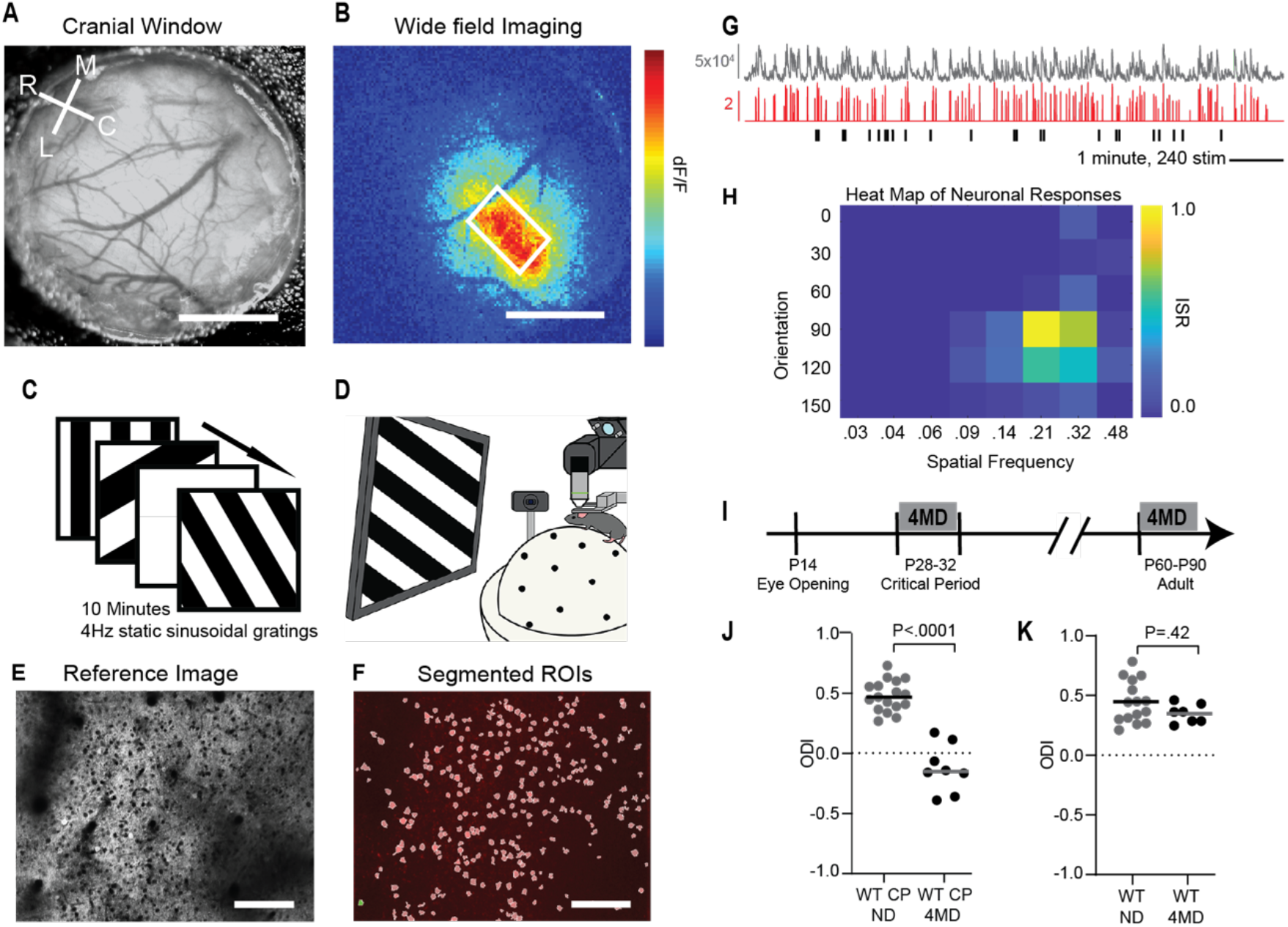
Calcium imaging at neuronal resolution detects OD plasticity confined to the developmental critical period. (**A**) An example cranial window 3mm in diameter centered over visual cortex. Upper left corner, coordinate axes. Scale bar = 1mm. (**B**) Wide field calcium imaging of neural activity in response to a horizontal bar 30 degrees wide and 2 degrees high drifting top to bottom at 10 degrees per second. White square encapsulates imaging field presented in panels E and F. Scale bar = 1mm. (**C**) The visual stimulus is sinusoidal gratings (not square as depicted) at 30 degrees intervals of orientation and between 0.028 and 0.48 cycled per degree in spatial frequency (SF) spaced at half octaves. In addition, an iso-luminant grey screen is presented. Stimuli are presented randomly at 4 Hz for 10 minutes. Each combination of orientation and SF is presented 40 times on average (range 29–56). (**D**) Illustration of calcium imaging setup. Alert mice are head-fixed freely running on a spherical treadmill floating on a column of air and positioned with monitor 35cm away centered on azimuth at zero elevation. An occluder (not depicted) is placed in front of contralateral or ipsilateral eye in separate imaging sessions. A camera records pupil diameter. (**E**) Example image of visual cortex recorded during an experiment. Imaging field is 750μm x 500μm. Scale bar = 100μm. (**F**) White circles correspond to manually identified ROIs for panel E. The ROI colored green (*lower left corner*) is presented in panel G. Scale bar = 100μm. (**G**) A representative trace of fluorescence over time for an ROI (*grey, top*) and the corresponding inferred spike rate (ISR) (*red, middle*). The timing of presentations of the preferred stimuli (90 deg, 0.21 cpd) are represented by vertical lines (*black, bottom*). Scale bar = 1 minute and 240 stimuli. (**H**) Heat map of the ISR for all combinations of orientation and SF. (**I**) Timeline of imaging and monocular deprivation. (**J**) Average ODI for critical period mice with and without 4 days of monocular deprivation (WT CP ND, mean ODI = 0.48, n = 17 mice; WT CP 4MD P32 mice, mean ODI = −0.12, n = 8 mice; one-way ANOVA with Sidak’s multiple comparisons test). Experimental groups augmented from Brown and McGee, 2023. (**K**) Average ODI for mice after closure of critical period with and without 4 days of monocular deprivation (WT ND mean ODI = 0.45, n = 15 mice; WT 4MD mice, mean ODI = 0.35, n = 7 mice, one-way ANOVA with Sidak’s multiple comparisons test).

Next, we leveraged the phenotype of mice lacking a functional gene for *nogo-66 receptor 1* (*ngr1*) to test how instability of neuronal tuning relates to the capacity for OD plasticity. The critical period does not close properly in *ngr1* mutants as they retain OD plasticity in response to brief MD as adults (Frantz et al. 2020; McGee et al. 2005; Stephany et al. 2014, 2016). The OD plasticity displayed by adult constitutive *ngr1* ‘knock-out’ mice (KO) is indistinguishable from that exhibited by WT mice during the critical period, including the time course, magnitude, laminar progression, sensitivity to barbiturates, and sensitivity to benzodiazepines (Fischer et al. 2007; Frantz et al. 2020; McGee et al. 2005; Pham et al. 2004; Stephany et al. 2016). Adult *ngr1* KO mice also display the intracortical disinhibition with 1-2d of MD that is the first known circuit adaptation following MD during the critical period (Kuhlman et al. 2013; Stephany et al. 2016). Thus, comparing the neuronal tuning of adult WT and *ngr1* KO mice permits isolating the circuit features associated with OD plasticity from other forms of plasticity that may be coincident but unrelated to this critical period during development.

Adult *ngr1* KO mice display OD plasticity as measured by calcium imaging. Adult mice (P60-90) receiving 4 days of MD possess lower mean ODI values than non-deprived mice (Fig. 2A). In the mouse, OD plasticity during the critical period is driven by a depression of aggregate neuronal responses to visual stimuli presented to the eye receiving MD (Frenkel & Bear 2004; Sato & Stryker 2008). At neuronal resolution, OD plasticity results from a decrease in the average response amplitude for neurons responding to visual stimuli presented to the contralateral eye, an increase in the percentage of neurons that are predominantly monocular neurons that only respond to visual stimuli presented to the ipsilateral eye (ODI = -1), a decrease in the percentage of predominantly contralateral monocular neurons (ODI = 1), and a modest reduction in the ODI values of binocular neurons (Brown & McGee 2023) (Fig. S2). Adult mice lacking *ngr1* display these same cellular mechanisms of OD plasticity (Fig. 2 B-C). In addition, MD alters the distribution of neurons for adult *ngr1* KO mice when plotted in as histograms of ODI or viewed a fields of neurons per mouse (Fig. 2 D-I).

**Fig 2.**
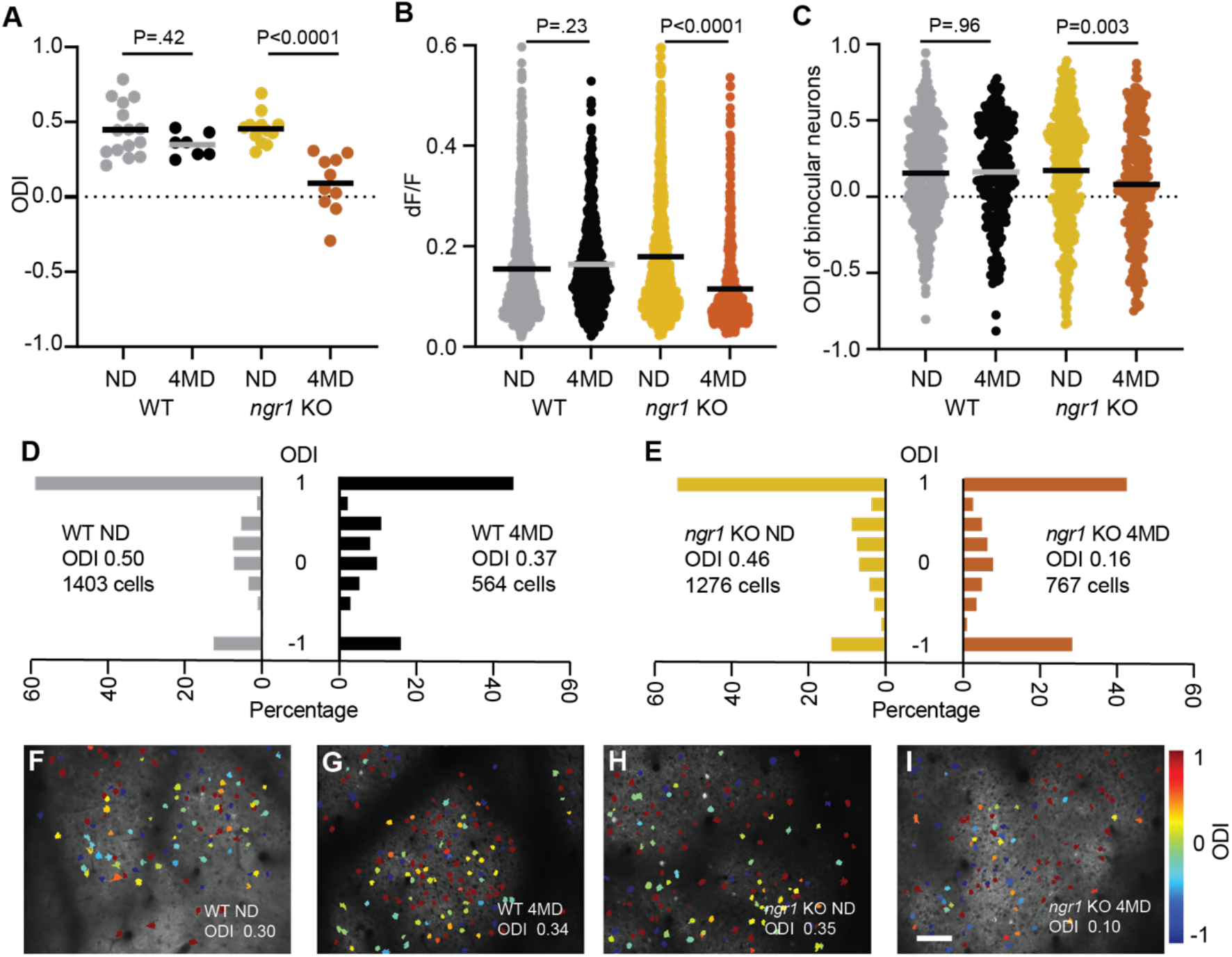
OD plasticity of adult *ngr1* KO mice at neuronal resolution. (**A**) Average ODI per mouse for adult WT and *ngr1* KO non-deprived (ND) mice and after 4 days of MD (4MD): (WT ND ODI = 0.45, n = 15 mice; WT 4MD mice, mean ODI = 0.35, n = 7 mice; *ngr1* KO mean ODI = 0.45, n =12 mice; *ngr1* KO 4MD mice mean = 0.09 n = 10 mice, one-way ANOVA with Sidak’s multiple comparisons test). Horizonal lines represent means. WT mice are presented from Fig. 1K for comparison (**B**) dF/F of contralateral neurons from WT ND, WT 4MD, *ngr1* KO ND, *ngr1* KO 4MD groups: (WT ND = 0.15, n= 1222 neurons; WT 4MD = 0.16, n = 475 neurons; *ngr1* KO ND = 0.17, n =1100 neurons; *ngr1* KO 4MD = 0.11 n = 550 neurons; one-way ANOVA Sidak’s multiple comparisons test). (**C**) ODI values of binocular neurons from WT ND, WT 4MD, *ngr1* KO ND, *ngr1* KO 4MD groups: (WT ND = 0.15, n= 391 neurons; WT 4MD = 0.16, n = 220 neurons; *ngr1* KO = 0.17, n =415 neurons; *ngr1* -/-4MD mice mean = 0.08, n = 226 neurons; one-way ANOVA with Sidak’s multiple comparisons test). (**D**) Histogram of neuronal ODI values for WT ND and 4MD groups: (WT ND mean ODI = 0.51, 1403 neurons; WT 4MD mean ODI = 0.36, 564 neurons). (**E**) Histogram of neuronal ODI values for *ngr1* KO ND and 4MD groups: (ngr1 KO mean ODI = 0.46, 1276 neurons; *ngr1* KO 4MD mean ODI = 0.17, 767 neurons). (**F-I**) Example fields of neurons for WT ND, WT 4MD, *ngr1* KO ND, *ngr1* KO 4MD mice. Neurons are color coded by ODI. Red neurons respond to contralateral eye. Blue neurons respond to the ipsilateral eye. Other colors respond to stimuli from both eyes. The mean ODI for the imaging field is stated in the lower right corner. Scale bar = 100μm.

Then, we applied this experimental paradigm to track the tuning properties of excitatory neurons in L2/3 of visual cortex across an interval of 4 days for WT and *ngr1* KO mice. We calculated binocularity hundreds of neurons for juvenile non-deprived WT mice, adult non-deprived WT mice, adult non-deprived *ngr1* KO mice, and adult *ngr1* KO mice receiving MD after the first imaging time point (Fig. 3). In addition to neurons that were consistently visually responsive, some neurons that were not visually responsive during the first imaging session (NR) became responsive to visual stimuli, while others which were previously visually responsive became non-responsive. The overall distribution of binocularity did not change between time points for non-deprived juvenile WT mice, adult WT mice, or adult *ngr1* KO mice as the ratios of predominantly contralateral monocular neurons (C), binocular neurons (B), and predominantly ipsilateral monocular neurons (I), were quite similar at both time points (Fig. 3 A-C). In contrast, adult *ngr1* KO mice receiving 4 days of MD displayed OD plasticity that reduced the percentage of C neurons and increased the percentage of I neurons, consistent with the results from the single time-point calcium imaging experiments (Fig. 2).

**Fig 3.**
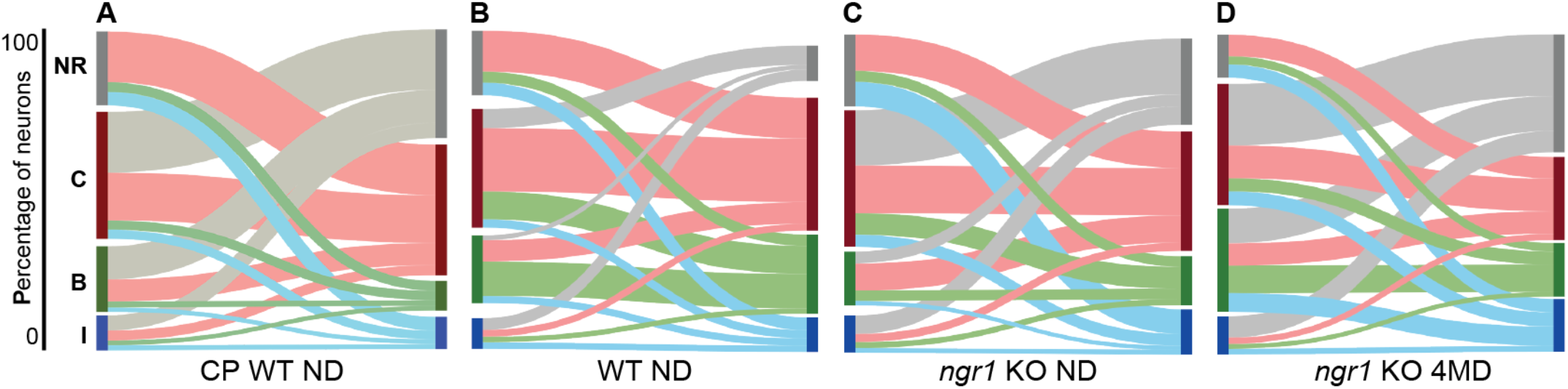
Reorganization of tuning for binocularity for longitudinally tracked neurons. (**A**) Sankey plots of ODI values for neurons imaged at P28 and again at P32 for in CP ND animals. Vertical bars on left side represent neuronal ODI at first imaging session. The length of the bar is proportional to the size of the population with that tuning preference. The grey bar represents non-visually evoked neurons (NR), the red bar represents neurons predominantly contralateral (**C**), the green bar represents neurons are binocular (**B**), and the blue bar represents predominantly ipsilateral (**I**) neurons. The lines connect to the tuning properties of neurons at second imaging session for days later (n =494 neurons, n=5 mice). The thickness of the line is proportional to the size of the population. (**B-D**) Sankey plots of ODI values of adult WT ND (n =780 neurons, n=6 mice), adult *ngr1* KO ND mice (n =595 neurons, n=4 animals), and adult *ngr1* KO 4MD mice (n =1062 neurons, n=7 animals).

However, there was considerable reorganization of the tuning preferences of individual neurons (C,B, or I) despite the conserved overall distribution of binocularity across the population of visually-responsive neurons for non-deprived mice (Fig. 3). While many neurons maintained stable tuning binocularity at both imaging time points (stable), other neurons were plastic and possessed different tuning (plastic). We calculated the percentage of stable and plastic neurons for these four groups (Fig. 4A). The percentage of plastic neurons with tuning instability was greater in juvenile WT mice than adult WT mice. This finding is consistent with a prior study measuring the stability of tuning for exclusively binocular neurons (B) in L2/3 of mouse V1 during the critical period and in adulthood (Tan et al. 2020). Interestingly, adult *ngr1* KO mice exhibited a greater percentage of plastic neurons than adult WT mice which was similar to that of juvenile WT mice.

**Fig 4.**
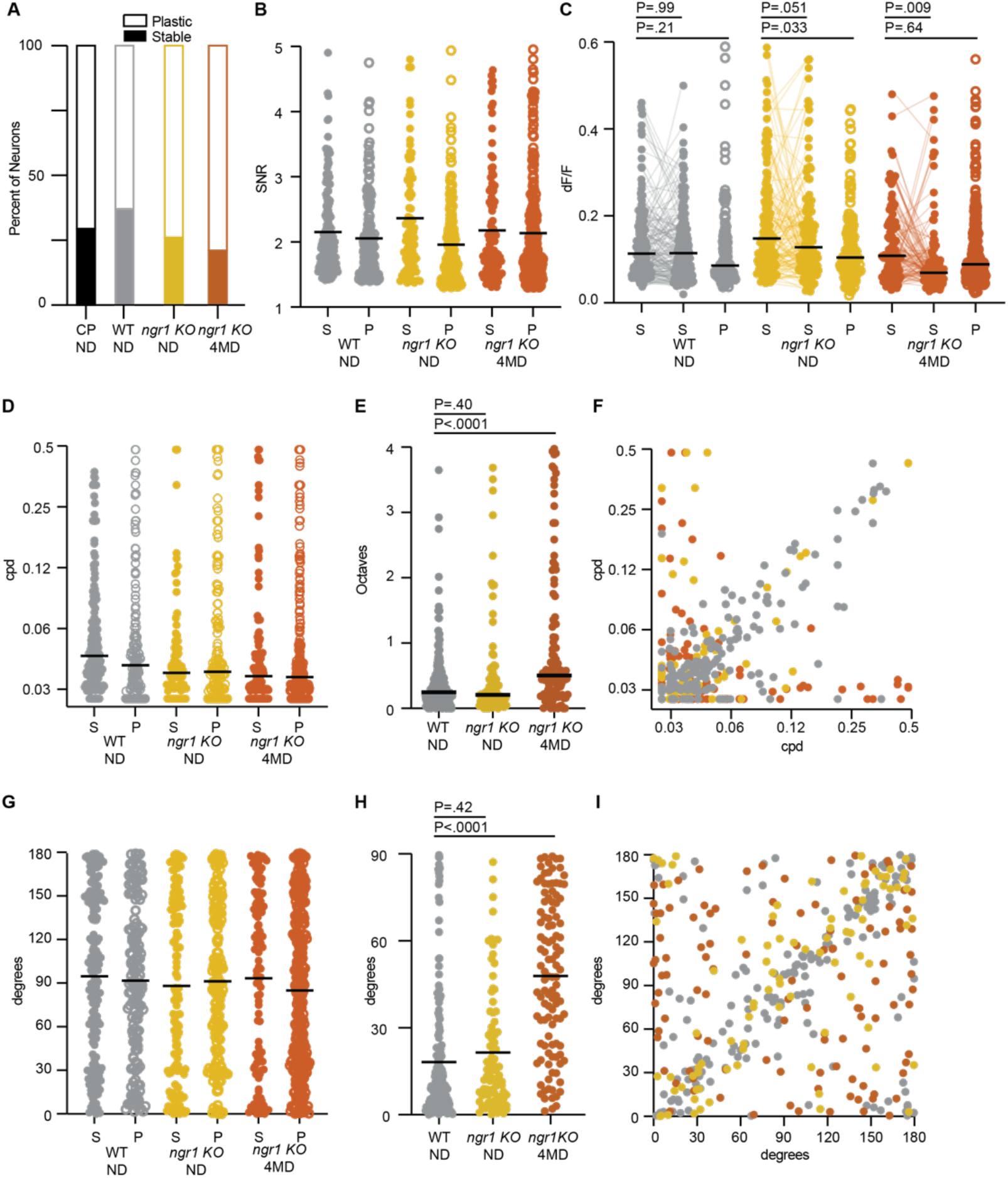
Response properties of longitudinally imaged neurons. (**A**) Bar graph depicting proportion of neurons with stable tuning (S) verses instable tuning (plastic, P) for binocularity from CP WT ND, WT ND, *ngr1* KO ND, *ngr1* KO 4MD groups (CP ND n = 494; WT ND =780; *ngr1* KO ND n =595; *ngr1* KO 4MD n =1062 total). (**B-I**) Tuning properties for stable and plastic predominantly contralateral neurons from WT ND (S=175, P=153), *ngr1* KO ND (S=87, P=164), *ngr1* KO 4MD (S=110, P=298) groups from panel A. (**B**) Signal:noise ratio (SNR): (WT ND S = 2.15, P = 2.05; *ngr1* KO ND 2.36, P = 1.95; *ngr1* KO 4MD S = 2.18, P = 2.12). Horizonal lines represent means (**C**) dF/F: (WT ND S1 = 0.14, S2 = 0.13, P = 0.12; *ngr1* KO ND S1 = 0.18, S2 = 0.14, P = 0.13; *ngr1* KO 4MD S1 = 0.13, S2 = 0.10, P = 0.12; Mixed effects analysis for repeated measures and Brown-Forsythe and Welch ANOVA). Lines connect individual stable neurons. Horizonal lines represent means. (**D**) Spatial frequency (SF) preference: (WT ND S = 0.045, P = 0.041; *ngr1* KO ND S = 0.037, P = 0.038; *ngr1* KO 4MD S = 0.036, P = 0.036). Horizonal lines represent medians. (**E**) Change in SF preference measured in octaves for stable neurons between time points (WT ND = 0.25; *ngr1* KO ND = 0.21; *ngr1* KO 4MD = 0.50; Brown-Forsythe and Welch ANOVA with Dunnett’s multiple comparisons test). Horizonal lines represent medians. (**F**) Scatter plot for preferred SF for stable neurons at both time points presented in panel E. (G) Orientation preference in degrees: (WT ND S = 92, P = 92; *ngr1* KO ND S = 90, P = 92; *ngr1* KO 4MD S = 94, P = 85). Horizonal lines represent medians. (**H**) Change in preferred orientation measured in degrees for stable neurons between time points (WT ND = 18; *ngr1* KO ND = 21; *ngr1* KO 4MD = 48; Brown-Forsythe and Welch ANOVA with Dunnett’s multiple comparisons test). Horizonal lines represent means. (**I**) Scatter plot for preferred Orientation for stable neurons at both time points presented in panel H.

Last, we explored the features of neurons with tuning instability within the response strength and tuning properties of these neuronal populations. We formulated several discrete hypotheses before analyzing the data: 1) tuning instability is an artefact of our inclusion criteria for visual responsiveness; 2) our measurements lack sensitivity to detect meaningful differences; and 3) plastic neurons represent only a subset of high spatial frequency (SF) or orientation preference. We confined our analysis to predominantly contralateral monocular neurons (C) because this was the population with the best sampling.

The inclusion criteria we employed to discriminate visually-responsive neurons from non-responsive neurons is an intersectional method that thresholds based on the ratio of the number of responses relative to presentations (spike ratio), and the ratio of signal:noise (SNR), for the visual stimulus encompassing the preferred SF and orientation for each neuron (Brown et al. 2024; Brown & McGee 2023). These criteria overlap with those applied by other labs measuring these tuning properties for excitatory neurons in mouse visual cortex with calcium imaging using drifting gratings and the same calcium sensor GCaMP6s (Jeon et al. 2018; Tan et al. 2020, 2021). If tuning instability were a consequence of neurons straddling the inclusion criteria for visual responsiveness, then they would be clustered near these threshold values. However, plastic neurons with tuning instability for binocularity (plastic) displayed similar mean spike ratio and SNR to neurons with stable tuning (Fig. 4B and S3). Critically, these plastic neurons, irrespective of genotype or deprivation, spanned the same range of magnitude as stable neurons. These findings are not consistent with the hypothesis that tuning instability is an artefact of our inclusion criteria.

To assess the sensitivity our measurements, we compared the average response magnitude to the preferred stimuli (dF/F) for neurons with stable tuning to neurons with tuning instability. Again, the distribution of the dF/F values were similar across genotypes and deprivation (Fig. 4C). Plastic neurons displayed slightly lower average dF/F, but the effect size was modest. In contrast, stable neurons from *ngr1* KO mice receiving 4 days of MD displayed the reduced dF/F amplitudes associated with critical-period OD plasticity, confirming the sensitivity of these measurements for detecting significant differences (Brown & McGee 2023). Neurons with tuning instability for binocularity also possessed the same range of preferred SF and orientation, although MD increased the average magnitude of difference in preferred SF and orientation between time points for stable neurons in adult *ngr1* mutant mice (Fig. 4D-I). Thus, the instability of tuning cannot be attributed to neurons with weaker responses nor predicted from different tuning parameters.

In summary, we conclude that the tuning of significant fraction of excitatory neurons in L2/3 of mouse visual cortex is unstable. The fraction of neurons with tuning instability is larger during the critical period, as well as in adult mice lacking a gene required to close the critical period. In the presence of consistent visual experience, the magnitude of tuning instability is not evident as the distribution of tuning properties is unaltered at the population level despite the interchange of tuning properties by individual neurons. Yet, perturbing vision with MD reveals OD plasticity in mice with greater tuning instability. We propose that experience-dependent plasticity adapts populations of neurons to recent experience by directing the instability of neuronal tuning to reorganize the distribution of tuning properties and thereby the function neural circuits.

What distinguishes cortical neurons with tuning instability from neurons with more stable tuning remains unclear. Neurons with tuning instability for binocularity cannot be predicted from response strength or the stability of tuning for SF or orientation. Whether this phenomenon is best described a ‘representation drift’ is unclear because distribution of tuning properties is conserved over time; there is no evident aggregate ‘drift’ of tuning properties without disruptive experience. Perhaps a transcriptional signature or signaling pathway regulates the stability of neuronal tuning. Identifying the molecular profile of tuning instability may benefit from a better understanding of the function of ngr1, as well as whether other strategies for promoting experience-dependent plasticity also increase tuning instability, including removing perineuronal nets (Pizzorusso et al. 2002), altering cholinergic tone (Morishita et al. 2010), treatment with psychedelics (Nardou et al. 2023), or reducing myelination (Xin et al. 2024).

## Methods

**Table 1.** Reagents and resources employed in the study

| REAGENT or RESOURCE | SOURCE | IDENTIFIER |
| --- | --- | --- |
| **Deposited data** |  |  |
| Calculated tuning properties for all neurons |  | N/A |
| **Experimental models: Organisms/strains** |  |  |
| Mouse: B6;DBA-Tg(tedO-GCaMP6s)2Niell/j | The Jackson Laboratory | RRID: ISMR_JAX: 024742 |
| Mouse: B6;Cg-Tg(Camk2a-tTA)1Mmay/DboJ | The Jackson Laboratory | RRID: ISMR_JAX: 007004 |
| Mouse: nogo-66 receptor (ngr1/rtn4r) constitutive mutant mice |  | Kim et al, 2004 |
| **Software and algorithms** |  |  |
| MATLAB | Mathworks | https://www.mathworks.com/ |
| Processing2 | Processing | https://processing.org/ |

### Lead contact

Further information and requests for resources and reagents should be directed to and will be fulfilled by the Lead Contact, Aaron W. McGee.

### Materials availability

This study did not generate new unique reagents

### Experimental model and subject details

All procedures were approved by University of Louisville Institutional Animal Care and Use Committee (IACUC) protocol 22105 and were in accord with guidelines set by the US National Institutes of Health. Mice were anesthetized by isoflurane inhalation and killed by carbon dioxide asphyxiation or cervical dislocation following deep anesthesia in accordance with approved protocols. Mice were housed in groups of 5 or fewer per cage in a 12/12 light–dark cycle. Animals were naive subjects with no prior history of participation in research studies. A total of 85 mice, both male (44) and female (41) were used in this study. The following male and female mice are represented in the following groups: one time point calcium imaging, P28-32 non-deprived WT, 9 males and 8 females; P32 4MD WT, 4 males and 4 females; P60-90 non-deprived WT, 7 males and 8 females; P60-90 4MD WT, 4 males and 3 females; P60-90 non-deprived *ngr1* KO, 6 males and 6 females; P60-90 4MD *ngr1* KO, 5 males and 5 females; two time point repeat calcium imaging, P60-90 non-deprived WT, 4 males and 2 females; P60-90 non-deprived *ngr1* KO repeat imaging, 2 males and 2 females; P60-90 4MD *ngr1* KO, 3 males and 4 females.

### Mice

Imaging was performed on double transgenic mice expressing GCaMP6s in forebrain excitatory neurons. The CaMKII-tTA (stock no. 007004) and TRE-GCaMP6s (stock no. 024742) transgenic mouse lines were obtained from Jackson Labs (Mayford et al. 1996; Wekselblatt et al. 2016) Mice were genotyped with primer sets suggested by Jackson Labs. These mice were crossed onto the *ngr1* KO background (Kim et al. 2004).

### Cranial windows

Wide field epi-fluorescent calcium imaging and two-photon calcium imaging were performed though a cranial window (Trachtenberg et al. 2002). In brief, mice were administered carprofen (5 mg/kg) and buprenorphine (0.1 mg/kg) for analgesia and anesthetized with isoflurane (4% induction, 1% to 2% maintenance). The scalp was shaved and mice were mounted on a stereotaxic frame with palate bar and their body temperature maintained at 37°C with a heat pad controlled by feedback from a rectal thermometer (TCAT-2LV, Physitemp). The scalp was resected, the connective tissue removed from the skull, and a custom aluminum headbar affixed with C&B metabond (Parkell). A circular region of bone 3 mm in diameter centered over left visual cortex was removed using a high-speed drill (Foredom). Care was taken to not perturb the dura. A sterile 3 mm circular glass coverslip was sealed to the surrounding skull with cyanoacrylate (Pacer Technology) and dental acrylic (Ortho-ject, Lang Dental). The remaining exposed skull likewise sealed with cyanoacrylate and dental acrylic. Mice recovered on a heating pad. Mice were left to recover for at least 2 days prior to 2-photon imaging.

### Wide field epi-fluorescent calcium imaging

After implantation of the cranial window and before 2-photon imaging, the binocular zone of visual cortex was identified with wide field calcium imaging similar to our method for optical imaging of intrinsic signals (Frantz et al. 2016; Kalatsky et al. 2005). In brief, mice were anesthetized with isoflurane (4% induction), provided a low dose of the sedative chlorprothixene (0.5 mg/kg IP; C1761, Sigma) and secured by the aluminum headbar. The eyes were lubricated with a thin layer of ophthalmic ointment (Puralube, Dechra Pharmaceuticals). Body temperature was maintained at 37°C with heating pad regulated by a rectal thermometer (TCAT-2LV, Physitemp). Visual stimulus was provided through custom-written software (MATLAB, Mathworks). A monitor was placed 25 cm directly in front of the animal and subtended +40 to −40 degrees of visual space in the vertical axis. A horizonal white bar (2 degrees high and 20 degrees wide) centered on the zero-degree azimuth drifted from the top to bottom of the monitor with a period of 8 seconds. The stimulus was repeated 60 times. Cortex was illuminated with blue light (475 ± 30 nm) (475/35, Semrock) from a stable light source (intralux dc-1100, Volpi). Fluorescence was captured utilizing a green filter (HQ620/20) attached to a tandem lens (50 mm lens, Computar) and camera (Manta G-1236B, Allied Vision). The imaging plane was defocused to approximately 200 μm below the pia. Images were captured at 10 Hz as images of 1,024 × 1,024 pixels and 12-bit depth. Images were binned spatially 4 × 4 before the magnitude of the response at the stimulus frequency (0.125 Hz) was measured by Fourier analysis.

### Visual stimuli and two-photon calcium imaging

Visual stimulus presentation and image acquisition were both performed according to our published methods which were modified from published studies (Brown et al. 2024; Brown & McGee 2023; Jimenez et al. 2018; Tan et al. 2020). In brief, a battery of static sinusoidal gratings was generated in real time with custom software (Processing, MATLAB). Stimulus presentation was synchronized to the imaging data by time stamping the presentation of each visual stimulus to the image acquisition frame number a transistor–transistor logic (TTL) pulse generated with an Arduino at each stimulus transition. Orientation was sampled at equal intervals of 30 degrees from 0 to 150 degrees (6 orientations). SF was sampled in 8 steps on a logarithmic scale at half-octaves from 0.028 to 0.48 cpd. An iso-luminant grey screen was included (blank) was provided as a ninth step in the SF sampling as a control. Spatial phase was equally sampled at 45-degree intervals from 0 to 315 degrees for each combination of orientation and SF. Gratings with random combinations of orientation, SF, and spatial phase were presented at a rate of 4 Hz on a monitor with a refresh rate of 60Hz. Imaging sessions were 10 minutes (2,400 gratings presented in total). Consequently, each combination of orientation and SF was presented 40 times on average (range 29 to 56). The monitor was centered on the zero azimuth and elevation 35 cm away from the mouse and subtended 45 (vertical) by 80 degrees (horizontal) of visual space.

Imaging was performed with a resonant scanning 2-photon microscope controlled by Scanbox image acquisition and analysis software (Neurolabware). The objective lens was fixed at vertical for all experiments. Fluorescence excitation was provided by a tunable wavelength infrared laser (Ultra II, Coherent) at 920 nm. Images were collected through a 16× water-immersion objected (Nikon, 0.8 NA). Images (512 × 796 pixels, 520 × 740 μm) were captured at 15.5 Hz at depths between 150 and 400 μm. Eye movements and changes in pupil size were recorded using a Dalsa Genie M1280 camera (Teledyne Dalsa) fitted with 50 mm 1.8 lens (Computar) and 800 nm long-pass filter (Edmunds Optics). Imaging was performed on alert mice positioned on a spherical treadmill by the aluminum head bar affixed to the skull. The visual stimulus was presented to each eye separately by covering the fellow eye with a small custom occluder.

### Image Processing

Image processing was performed as described previously (Brown & McGee 2023; Tan et al. 2020). Imaging series for each eye were motion corrected with the SbxAlign tool. Regions of interest (ROIs) corresponding to excitatory neurons were selected manually with the SbxSegment tool following computation of pixel-wise correlation of fluorescence changes over time from 350 evenly spaced frames (∼4%). ROIs for each experiment were determined by correlated pixels the size similar to that of a neuronal soma. The fluorescence signal for each ROI and the surrounding neuropil were extracted from this segmentation map.

### Image Analysis

Image analysis was performed as described previously with minor modifications (Brown & McGee 2023). The fluorescence signal for each neuron was extracted by computing the mean of the calcium fluorescence within each ROI and subtracting the median fluorescence from the surrounding perimeter of neuropil (Ringach et al. 2016; Tan et al. 2020). An inferred spike rate (ISR) was estimated from adjusted fluorescence signal with the Vanilla algorithm (Berens et al. 2018). A reverse correlation of the ISR to stimulus onset was used to calculate the preferred stimuli (Brown & McGee 2023; Jimenez et al. 2018; Ringach et al. 2016; Tan et al. 2020). Neurons that satisfied 3 criteria were categorized as visually responsive: (1) the ISR was highest with the optimal delay of 4 to 9 frames following stimulus onset. This delay was determined empirically for this transgenic GCaMP6s mouse (Brown & McGee 2023; Tan et al. 2020); (2) the SNR was greater than 1.3. The signal is the mean of the spiking standard deviation at the optical delay between 4 and 9 frames after stimulus onset and the noise this value at frames −2 to 0 before the stimulus onset or 15 to 18 after it (Jimenez et al. 2018; Tan et al. 2020). (3) and neuron responded to at least 13% of the presentations of the preferred stimulus. Visual responsiveness for every neuron was determined independently for each eye. The visual stimulus capturing the preferred orientation and SF was the determined from the matrix of all orientations and SFs presented as the combination with highest average ISR.

The preferred orientation for each neuron was calculated as:

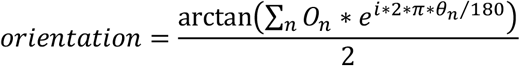

On is a 1 × 6 array of the mean z-scores associated with the calculation of the ISR at orientations Θn (0 to 150 degrees, spaced every 30 degrees). Orientation calculated with this formula is in radians and was converted to degrees. The tuning width was the full width at half-maximum of the preferred orientation.

The preferred SF for each neuron was calculated as:

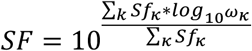

Sfk is a 1 × 8 array of the mean z-scores at SFs ωk (8 equal steps on a logarithmic scale from 0.028 to 0.481 cpd). Tails of the distribution were clipped at 25% of the peak response. The tuning width was the full width at half-maximum of the preferred SF in octaves. The percent visually responsive neurons with significant responses at each SF was determined by comparing the distribution of ISR values at each SF versus the stimulus blank with a KW-test with Dunn’s correction for 8 comparisons. Neurons with P < 0.01 for a given SF were considered significant responses at that SF (Salinas et al. 2017).

### Ocular Dominance Index (ODI)

Neuronal ODI was calculated as (C-I)/(C+I), where C and I are the mean normalized change in fluorescence (dF/F) for the preferred visual stimulus for the contralateral eye and ipsilateral eye, respectively. In cases where neurons displayed no significant response to visual stimuli provided to one eye, they were considered monocular for the other eye and assigned ODI values of 1 (contralateral) and -1 (ipsilateral) Neuronal ODI was calculated as (C − I) / (C + I), where C and I are the mean normalized change in fluorescence (dF/F) for the preferred visual stimulus for the contralateral eye and ipsilateral eye, respectively. In cases where neurons displayed no significant response to visual stimuli provided to one eye, they were considered monocular for the other eye and assigned ODI values of 1 (contralateral) and −1 (ipsilateral) (Salinas et al. 2017).

Summed ODI was calculated by summing the dF/F for the preferred visual stimulus for all neurons visually responsive to the contralateral eye (C) and ipsilateral eye (I) for each mouse, respectively. The summed ODI per mouse was then calculated as (C − I) / (C + I) for each mouse.

### Repeated two-photon calcium imaging

Each imaging session was segmented independently, and every ROI was assigned a unique number. No geometric transformations were performed to match segmentation masks for ROIs from the 2 imaging sessions. The segmentation masks for the 2 imaging sessions were then compared and ROIs with at least 50% overlap were considered the same neuron. A perimeter of neurons with overlapping ROIs and tuning properties that did not change between imaging sessions (a difference in orientation preference of less than 30 degrees and SF preference of less than an octave) defined the matched imaging plane. To determine the SNR values of lost neurons at P32 and gained neurons at P28, the segmentation masks were exchanged between time points and the SNR from ROIs for the corresponding neurons at the other time point were calculated.

### Monocular deprivation by lid suture

The right eye was sutured closed with a single mattress suture with 6–0 Prolene monofilament (Ethicon 8709) (Stephany et al. 2014). Prior to imaging, mice were briefly (<5 minutes) anesthetized with isoflurane (4% induction, 1% to 2% maintenance), the suture removed with Vannas scissors (Fine Science Tools). The eye was flushed with sterile saline and examined for corneal abrasions with a stereomicroscope. The mouse was then immediately head-fixed for imaging and allowed to recover for no less than 45 minutes. The occluder was positioned over one eye as soon as the mouse was head-fixed and occluded one of the eyes at all times. At no point during the experiment were mice permitted unobstructed binocular vision.

### Statistics

No statistical methods were used to predetermine sample size. All statistical analyses were done using Prism 8 software (GraphPad Software). Multiple comparisons were tested with Analysis of Variance (ANOVA) tests and paired tests with Mixed-effects analysis.

## Acknowledgements

We thank D. Ringach (UCLA) and J. Trachtenberg (UCLA) for sharing software and hardware design for visual stimulus presentation and image analysis prior to publication, A. Eliasen and G. Armstrong for software development, and B. Croslin for mouse husbandry and genotyping. This research is supported by the National Eye Institute (R01EY035138 and R01EY35885 to AWM)

## Author Contributions

The contributions are as follows: TCB and AWM designed the study. TCB performed the experiments. TCB and AWM analyzed the data, built the figures, and wrote the manuscript.

