## Supplemental Figures for "Experience directs the instability of neuronal tuning for critical period plasticity in mouse visual cortex"

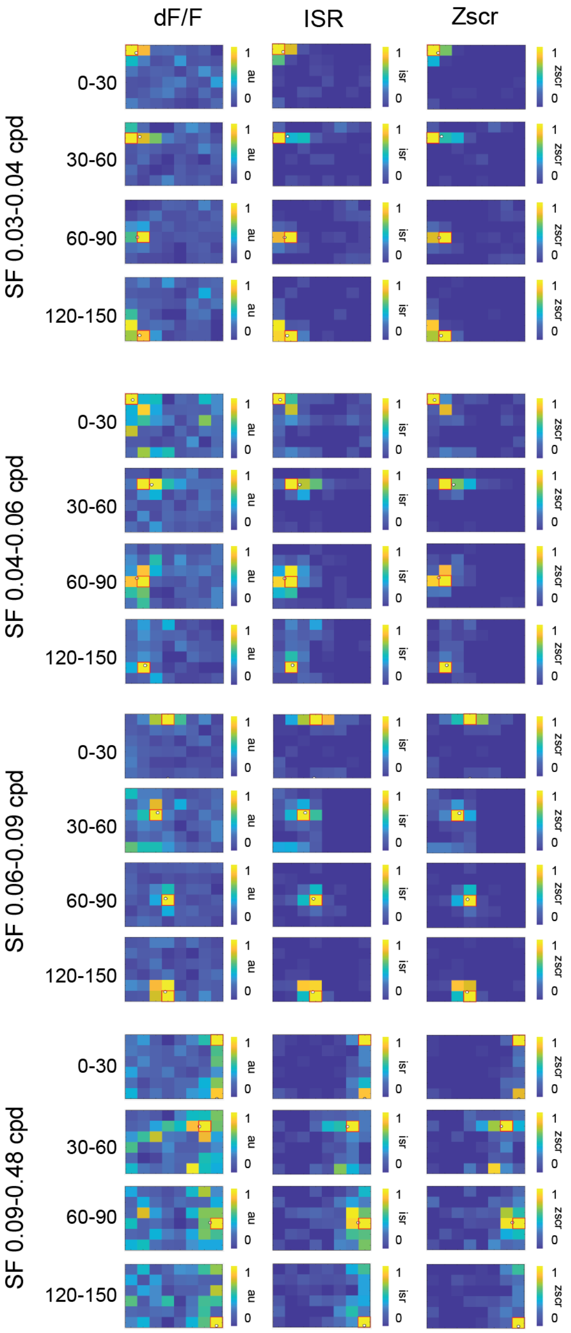


**Supplemental Fig. 1. Example heat maps of dF/F, ISR, and z-score for neurons spanning the range of orientations and spatial frequencies**
Example heat maps from representative neurons with tuning preferences across the range of presented spatial frequency and orientation. Each neuron is shown with a heat map generated for average change in fluorescence (dF/F), inferred spike rate (ISR), and Z score (Zscr). Heat maps are normalized to the maximum response.

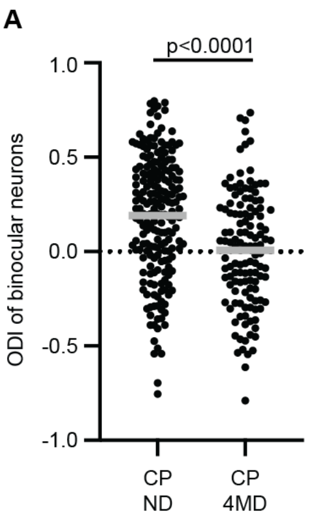


**Supplemental Fig. 2. ODI values of binocular neurons from juvenile non-deprived mice and after 4 days of MD.**
(**A**) Distribution of ODI values for binocular neurons from juvenile non-deprived mice and after 4 days of MD. Lines at means (CP ND mean = 0.19 n= 203 neurons; CP 4MD mice, mean = 0.01, n = 127 neurons, unpaired 2-way t-test witih Welch's correction). Other tuning properties for these groups published in Brown and McGee, 2023.

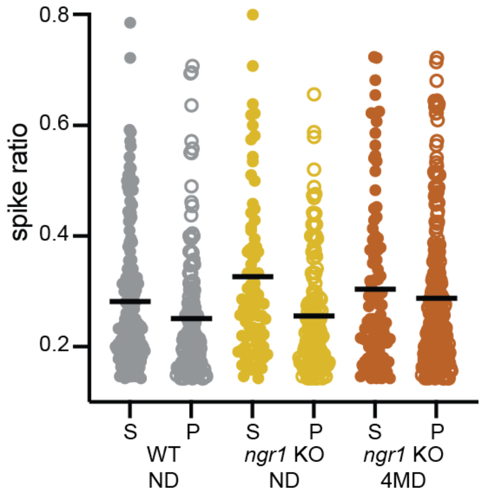


**Supplemental Fig. 3. Spike ratio of stable and plastic contralateral neurons for WT ND, *ngr1* KO ND, and *ngr1* KO 4MD groups.**
Plot of spike ratio for stable (S) and plastic (P) contralateral neurons: (WT ND S = 0.28, n =175 neurons, P = 0.25, n=153 neurons; *ngr1* KO ND S = 0.32, n =87 neurons, P = 0.24, n=164 neurons; ngr1 KO 4MD S = 0.29, n =110 neurons, P = 0.28, n=298 neurons). Horizontal bars indicate the mean.
